# Ecological and life history traits are associated with Ross River virus infection among sylvatic mammals in Australia

**DOI:** 10.1101/475814

**Authors:** Michael G. Walsh

## Abstract

**Background:** Ross River virus (RRV) is Australia’s most important arbovirus given its annual burden of disease and the relatively large number of Australians at risk for infection. This mosquito-borne arbovirus is also a zoonosis, making its epidemiology and infection ecology complex and cryptic. Our grasp of enzootic, epizootic, and zoonotic RRV transmission dynamics is imprecise largely due to a poor understanding of the role of wild mammalian hosts in the RRV system.

**Methods:** The current study applied a piecewise structural equation model (PSEM) toward an interspecific comparison of sylvatic Australian mammals to characterize the ecological and life history profile of species with a history of RRV infection relative to those species with no such history among all wild mammalian species surveyed for RRV infection. The effects of species traits were assessed through multiple causal pathways within the PSEM framework.

**Results:** Sylvatic mammalian species with a history of RRV infection tended to express dietary specialization and smaller population density. These species were also characterized by a longer gestation length.

**Conclusions:** This study provides the first interspecific comparison of wild mammals for RRV infection and identifies some potential targets for future wildlife surveys into the infection ecology of this important arbovirus. An applied RRV macroecology may prove invaluable to the epidemiological modeling of RRV epidemics across diverse sylvatic landscapes, as well as to the development of human and animal health surveillance systems.

## Background

Ross River virus (RRV) is simultaneously Australia’s most significant vector-borne and zoonotic pathogen. There are approximately 5100 cases annually[1], which is ten-fold higher than all other zoonoses[2]. Transmission of this alphavirus involves multiple mosquito vectors, which in turn exhibit heterogeneous host preferences in diverse dryland and wetland ecosystems[3]. Moreover, this heterogeneity modulates the dynamics of enzootic, epizootic and zoonotic RRV transmission, imparting a unique infection ecology to each. Furthermore, reservoir and amplification hosts among Australian wildlife have been shown to influence the landscape epidemiology of RRV, particularly with respect to human spillover[4, 5]. Nevertheless, the contribution of specific mammalian hosts to RRV epidemiology remains ill-defined despite several decades of human surveillance and wildlife sampling across Australia[1].

Ross River virus infection has been identified definitively in 21 native mammalian hosts in several field surveys and experimental studies[6–14]. However, many of these surveys target only single species in highly localized areas. None consider species’ ecology and how this may relate to 1) infection dynamics, such as the role of wildlife as maintenance or amplification hosts, or 2) the facilitation of human exposure in anthropogenically altered landscapes[15]. By contrast, identifying the biological and life-history characteristics of wildlife hosts has the potential to offer novel insight into the epidemiology and infection ecology of RRV and may suggest a specific wildlife-human interface as particularly vulnerable to spillover. This is a critical consideration as Australian sylvatic landscapes have undergone and continue to undergo rapid change due to habitat loss[16]. To date there has been no investigation of species-level biological and life history characteristics of RRV wildlife hosts and their comparison to non-hosts in endemic areas. Recent work in this domain has identified associations between species-level life history characteristics of wildlife reservoirs for other pathogens and human spillover[17]. This approach may prove useful in identifying traits associated with RRV infection status, thus enabling macroecology to inform sylvatic RRV epidemiology.

Recent studies have explored intriguing differences in life history characteristics related to the ways in which fast versus slow life history may mediate disease systems with high potential for spillover. Key delineators of fast living commonly include 1) high volume reproductive output with lower investment in early development as typified for example by short gestation, short inter-birth intervals, large litters, and early sexual maturity, and 2) a shorter lifespan. Conversely, slow living species would be characterized by lesser reproductive output with greater investment in early development as well as longer lifespans[18]. Some work has shown that fast life history is associated with greater reservoir competence[19, 20], and may in turn drive zoonotic transmission[20, 21], due to immunological mediation of transmission dynamics[22]. A comprehensive survey of all mammalian host-pathogen systems provided strong evidence confirming this relationship between reservoir hosts and living fast[17]. The current investigation explored whether the RRV system converged with, or diverged from, the typical fast life history of many mammalian host-pathogen systems that has been identified globally.

This study compared biological and life history traits of sylvatic mammalian species with demonstrated RRV infection to the traits of species without evidence of infection. Toward this end and as a causally sound approach to epidemiological inference, interspecific biological and life history characteristics were interrogated with respect to species’ RRV infection status using a piecewise structural equation model with phylogenetic generalized linear models as its component structures to account for the taxonomic correlation.

## Materials and methods

Forty-six sylvatic mammalian species surveyed for RRV infection were identified in the literature. Species reported as positive for infection by serology, polymerase chain reaction amplification, or isolation of virus were classified as RRV positive species [6–14, 23–27]. The remaining sylvatic mammal species sampled for RRV screening, but which tested negative, were classified as RRV negative species. It should be noted that the broad classification scheme based on the three modalities described above is not able to delineate RRV positive species as viral reservoirs or competent hosts. Rather, it simply marks these species as susceptible to infection. Twenty-one native mammalian species, comprising diprotodonts (11 macropods and 1 possum species), peramelemorphs (3 species), megabats (2 species), and rodents (4 species), were classified as RRV-positive, while 5 introduced mammalian species (brown rats, rabbits, pigs, dogs, and cats) were classified as positive. The full list of species classified as RRV hosts and non-hosts based on their documented infection status in the literature is presented in Additional file 1: Host list. Only wild/feral species were considered in this analysis because this study sought to describe the macroecology of sylvatic RRV infection history, which may offer unique insight into the epidemiological context of wilderness landscapes. Moreover, contextualizing zoonotic risk with macroecology may also present opportunities to integrate conservation and public health initiatives.

Species-level biological and life history characteristics were obtained from the PanTHERIA mammalian dataset[28]. The following life history traits were selected as representative metrics of the fast-slow continuum concept of life history[29], and which exhibited low levels of missing data (<70%) in the PanTHERIA database for the host taxonomies under study: gestation length (months), sexual maturity age (days), weaning age (months), litter size (number of animals per litter), inter-birth interval (months) and maximum longevity (months), i.e. lifespan; whereas adult body mass (kg), population density (animals/km^2^), home range (km^2^), and diet breadth were selected as additional ecologically relevant species traits that also exhibited low levels of missing data (Additional file 2). Species-level diet breadth describes the mean number of dietary categories consumed throughout the year, wherein food categories were classified as vertebrate, invertebrate, fruit, flowers/nectar/pollen, leaves/branches/bark, seeds, grass, and roots/tubers. Any remaining species-level missing data for these characteristics were imputed using a random forest machine learning algorithm, which has been shown to be a robust approach to this application[17, 30]. The algorithm was implemented using the `rfImpute` function in the randomForest package[31].

### Statistical Analysis

A piecewise structural equation model (PSEM) was used to identify associations between species’ infection history and species’ ecology and life history[32]. This approach is considered an optimal application to causal inference since it evaluates the covariance among all independent variables as part of the modeling process. Within the PSEM framework, relationships between covariates and between covariates and the outcome (RRV infection history) are estimated as individual structured equations to quantify and map direct and indirect associations. The implementation of local estimation rather than global estimation for these models also makes this approach more suited to small sample sizes. The PSEM thus comprises a set of equations quantifying causal paths in the system, and has become a popular approach to estimating causality in epidemiological relationships. Phylogenetic generalized linear models (PGLMs) were used as the set of equations for these causal paths in this analysis to account for the phylogenetic correlation [32]. To correct for potential bias introduced by differences in reporting effort across species, the number of individuals sampled per species surveyed (across all surveys) was used to quantify reporting effort and this was included as an additional covariate in these PGLM models[33, 34]. Furthermore, because of the relatively low number of species available with RRV infection history (n = 46), the maximum number of variables included in the final PGLM was constrained to ensure efficient parameter estimation and prevent overfitting. As such, 10 bivariate models were fitted wherein each model comprised a simple PGLM assessing the crude association between infection history and each species trait (although these crude models did adjust for reporting effort as described above): gestation length, sexual maturity age, weaning age, litter size, inter-birth interval, maximum longevity, population density, home range, diet breadth, and body mass. Using the Akaike information criterion, the four best-fitting trait variables were then selected and fitted together to assess the best fitting multivariable PGLM. Body mass was also included by default as this trait has the potential to modify many other ecological traits and is generally considered essential to account for when comparing species traits across orders[29].

The PSEM was then fitted to RRV infection status based on the best-fitting PGLM described above. Fisher’s C was used to test for conditional independence (p-value > 0.05 indicates there are no missing paths among variables) and the AIC was used to evaluate goodness-of-fit[35, 36]. The piecewiseSEM package[36] in the R statistical environment, v. 3.1.3[37] was used to implement the PSEM. The ape package was used to quantify the phylogenetic correlation[38, 39], and the phylolm package was to fit the phylogenetic generalized linear models used in the PSEM[40].

## Results

The final PGLM of RRV infection history and species traits is presented in Table 1. The individual bivariate PGLMs showing the goodness-of-fit of individual variables is presented in Additional file 3. The final PGLM model was a better fit (AIC = 87.9) than 1) a reduced model with just adult body mass, population density, and diet breadth (AIC = 95.0), 2) a reduced model with just gestation length and litter size (AIC = 99.8), or 3) models considering each life history and ecological trait sequentially (all AIC ≥ 94.6). Specific examination of the inclusion of species sample size as a proxy for reporting effort demonstrated this model to be a moderately poorer fit (AIC = 87.9) than the model excluding sample size (AIC = 86.5), so the latter was retained for the PSEM.

**Table 1.**
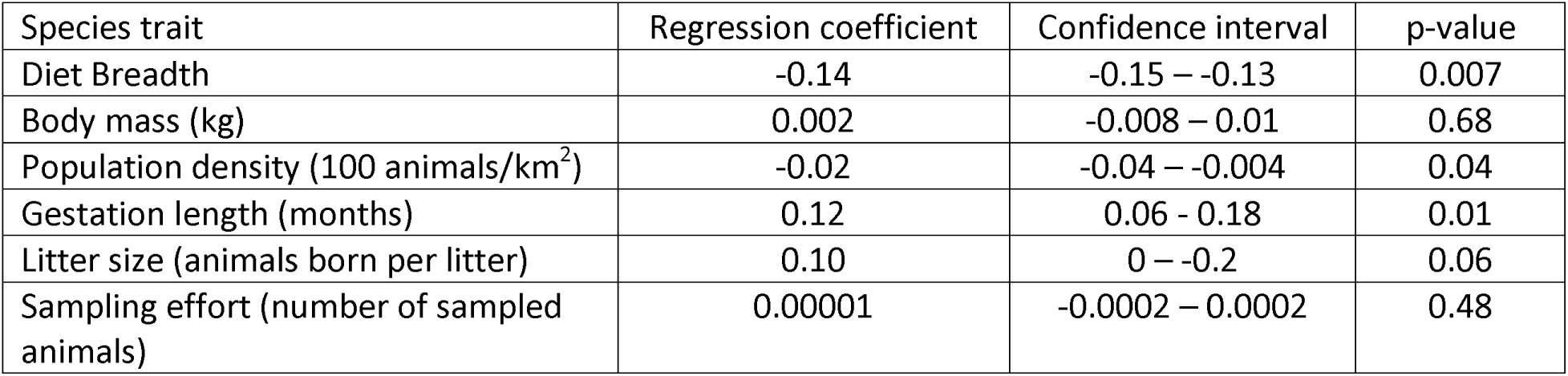
Phylogenetic generalized linear model of the associations between Ross River virus infection history and biological and life history traits among sylvatic Australian mammals.

Diet breadth (β = −0.14, 95% C.I. −0.15 – −0.13), population density (β = −0.02, 95% C.I. −0.04 – −0.004), and gestation length (β = 0.12, 95% C.I. 0.06 - 0.18) dominated the trait profile of RRV-susceptible species (Table 1), wherein increasing specialization and gestation length, and lower population density, were all associated with a positive infection history. The PSEM demonstrated that no additional associations with infection history were mediated through unanticipated indirect effects (Figure 1), suggesting that the single PGLM model adequately represented the relationships between species’ traits and infection history. The model that emerged among these covariates showed that diet breadth, population density, and gestation length were influential to RRV infection history among sylvatic mammals that have been surveyed for this arbovirus. Additionally, reporting effort was not associated with infection history, thus suggesting that RRV-positive species did not appear to be over-represented in sampling effort among the sylvatic mammals surveyed.

**Figure 1.**
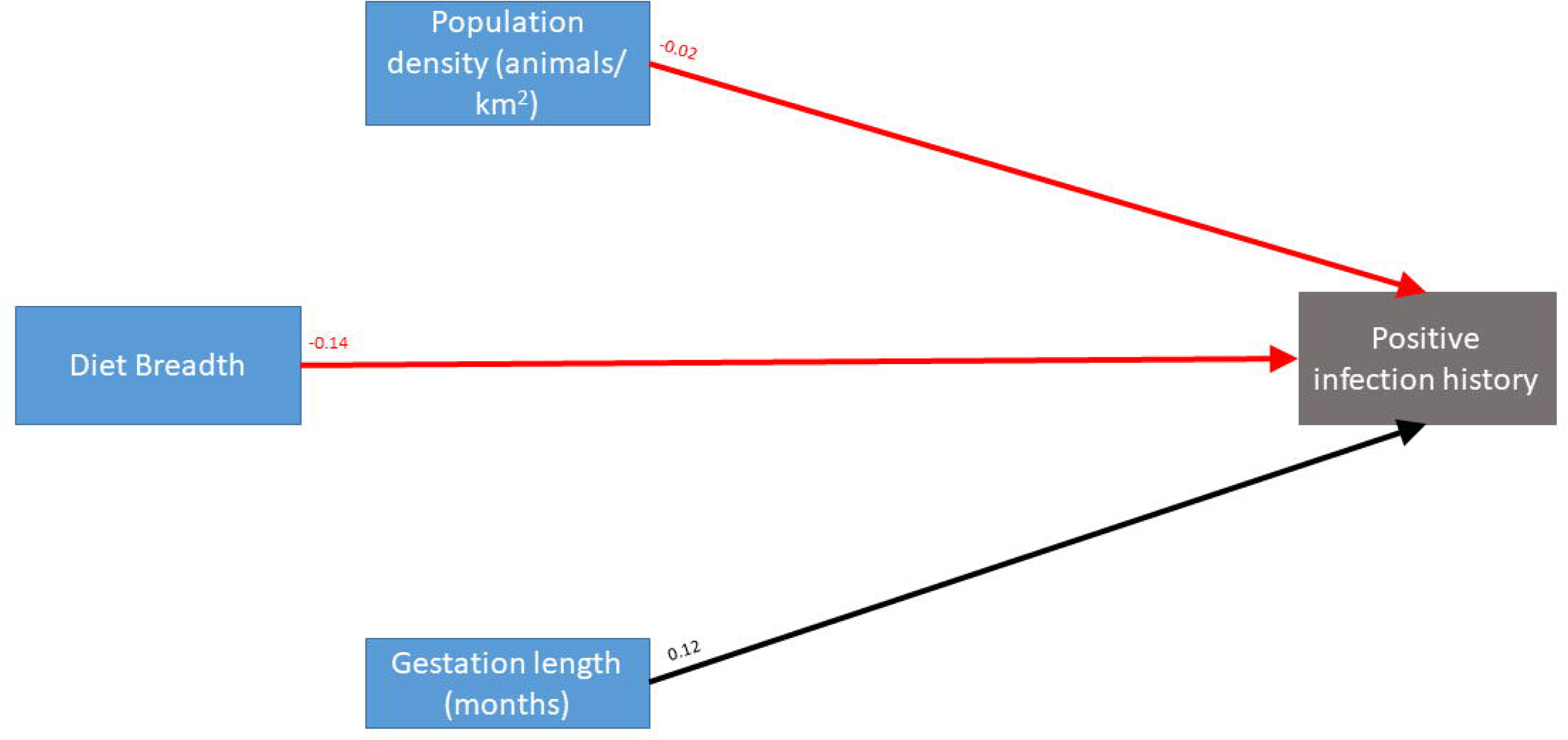
Piecewise structural equation model (PSEM), wherein nine structured equations were fit as individual components of the PSEM with phylogenetic generalized linear regression used to account for the correlation structure. The regression coefficients are presented above each path. Black lines indicate positive relationships between traits, while red lines indicate inverse relationships.

## Discussion

This investigation suggests that a narrower repertoire of food sources, indicating relative diet specialization, lower population density, and a tendency toward longer gestation characterize wild mammalian RRV infection history. The extent to which perturbations such has habitat loss or resource provisioning among competent hosts with this profile could enhance their potential to act as bridging species for zoonotic transmission is unknown from this, or previous, studies. Such phenomena were not testable in the current study because a clear delineation between competent and non-competent hosts was not possible. Nevertheless, this should be formally tested in future work through better virological surveillance of wildlife and by testing the epidemiological impact of habitat conservation on the prevention of zoonotic RRV transmission. Interestingly, slow-living, as indicated by longer gestation, emerged from the life history profile of RRV-positive species. However, this will also require validation with future work, particularly since only this single life history characteristic was relevant in the current study and the possibility that slow-living may simply reflect greater opportunity for exposure.

The ecological context for the relationships between RRV infection history and diet breadth and population density is not clear from the current study, however it may be that animals with a narrower dietary breadth may expend greater time in foraging, and possibly over a greater area, thereby exposing them to mosquito vectors for longer periods of time[41]. Despite its broader ecological significance, the relationship with dietary specialization is interesting because specialists also tend to be more sensitive to habitat fragmentation[42, 43]. While such changes can sometimes result in species’ inability to adapt to the altered landscapes, species with low diet breadth have also been shown to exhibit the lowest levels of population decline following habitat fragmentation[44]. As such, habitat fragmentation or resource provisioning may induce population perturbations that lead to the persistence of important maintenance hosts, or increase contact with these hosts, both of which in turn could enhance the potential for enzootic, epizootic and zoonotic transmission if biodiversity is also lost[45]. The inverse association with population is also interesting, but may seem counterintuitive given common that vector-borne diseases are commonly assumed to demonstrate frequency-dependent transmission dynamics[46]. It is worth noting, however that this characterization is generally made without adequate parameterization of local spatial heterogeneity in rates of contact, and frequency-dependence can mask density-dependence unless contact networks are adequately defined[47]. This association may reflect this phenomena, but it is also possible that the association with population density is an artifact of species with lower population density exhibiting greater dietary specialization[48], which exhibited a strong association with infection history in this study. As described above, the results from the current study are insufficient to make definitive claims given the inability to distinguish between competent and non-competent hosts, but rather suggest ecological contexts to explore in future RRV disease ecology fieldwork.

This study also contributes to a growing body of work endeavoring to define zoonotic infection ecology by classifying mammalian hosts according to species’ life history. The extent to which species “live fast” or “live slow” may have important implications for immunogenicity and pathogenicity[22], thereby influencing pathogen persistence and shedding and subsequent spillover to novel hosts[19–21]. This study provided some evidence that a slower life history, as characterized by longer gestation, is characteristic of RRV infection history among sylvatic mammals. It would appear that the RRV system in the Australian mammalian landscape is more complex with respect to life history than other host-pathogen systems that demonstrate a higher proclivity to infection susceptibility among fast-living mammals[17, 20, 21]. This contrast notwithstanding, a similar pattern of slow-living life history was identified in another vector-borne arbovirus with significant zoonotic disease burden[49]. The importance of increased early life investment may reflect the significance of species with fewer but extended pulses in reproductive outputs to the seasonal cycling of RRV transmission. For example, increased time to reach maturity may correspond to a longer period of immunological naivety, which may extend opportunities for infectious exposure to vectors to within the critical temporal window that defines the transmission dynamics of the seasonal cycling of RRV. Moreover, it has been shown that life history modulates immune function with fast-living species investing more in the development of innate immunity, while slow-living species invest more in adaptive immunity[50, 51], which may further affect the infection susceptibility of mammalian species[20–22]. Nevertheless, the association between infection history and slow life history may simply reflect a longer exposure opportunity for slow-life species. As the current study could not evaluate acute infection in these species, the findings require validation by studies that do.

The epidemiology of Australia’s most important vector-borne and zoonotic pathogen has heretofore been poorly informed by the infection ecology of its wildlife hosts. While macropods are typically considered the primary drivers of RRV circulation and spillover to humans[3, 7, 8], the current interspecific interrogation defines an ecological profile of potential wildlife hosts according to their biological and ecological traits, and thus expands the scope for more targeted modeling of zoonotic transmission[15]. Moreover, the current study helps to synthesize an ecological model from previous wildlife surveys, which have frequently been conducted in isolation. Of the wildlife surveys conducted to date, only five were generalist surveys that sampled a broad selection of potential wildlife species for evidence of RRV infection[7, 9, 11, 13, 26]. The remaining studies were either part of larger outbreak investigations or targeted single or limited species for study. A recent systematic review of RRV virology parameters in vertebrate hosts identified the highest seroprevalence surveyed in macropods and high viremia in experimentally infected marsupials[5]. However, while high in marsupials, this review also showed no significant difference in viremia between marsupials, placentals, and birds during experimental infection. Moreover, the investigators make a sound argument that placental hosts may also contribute to RRV circulation[5]. The current study largely agrees with this systematic review, highlighting the potential additional contribution of non-native animals to RRV transmission dynamics and therefore cautions that a broader focus may yet be required to definitively articulate enzootic, epizootic, and zoonotic transmission. Nevertheless, the current study focused specifically on wildlife to identify how macroecology may inform the epidemiology sylvatic RRV transmission. Wildlife species are an important focus in their own right given the rapid incursion of the human population into wildlife habitat and the largely unknown consequences of emerging wildlife-human interfaces on pathogen transmission dynamics. Wildlife species abundance is highly variable across diverse wetland and dryland ecosystems, is often sensitive to anthropogenic alteration of these landscapes, and can modify the presence, diversity, and frequency of vector mosquitoes, with further implications for zoonotic transmission risk[3, 52, 53].

Beyond what has already been described, this study has some additional limitations that are discussed in greater detail below. First, the number of wildlife species surveyed for RRV is relatively small (n = 46) and therefore some relationships may have been missed due to the diminished power of the sample. Second, as described in the Methods section, there were considerable missing data for some species’ biological and life history characteristics, which could not be included in these analyses. We imputed data where appropriate (i.e. variables within a threshold of <70% missingness) using methods previously employed[17]. As such, the imputed data represent a narrower spectrum of Australian mammal biology and life history traits, but it is expected that inferences based on these traits will be more reliable if also incomplete. Third, the classification of infection-positive versus infection-negative species was based on serology in many instances, which only identifies these animals as susceptible to infection and does not indicate definitively whether these are competent reservoir hosts that may serve to maintain, or amplify, virus circulation. Fourth, while the inferences made here with respect to RRV host traits are ecologically relevant, if the ultimate goal is a more complete description of zoonotic RRV epidemiology then this will require the landscape analysis of these species, their mosquito vectors, and the coincident human cases in real time as a broad application of virology-based RRV surveillance in rural, urban, and peri-urban space. Valuable initial efforts have been made to define the landscape epidemiology of RRV, but only at coarse spatial[4, 54] and temporal scale[55, 56], and based on limited surveillance data.

## Conclusions

The findings described here highlight dietary specialization, population density, and gestation length as characteristic of RRV infection in wildlife, which provides the first such interspecific comparison and offers a richer understanding of RRV infection ecology. This applied RRV macroecology may add substantive value to the epidemiological modeling of RRV epidemics across varied wetland and dryland habitat. However, the value added will depend entirely on the commitment to active monitoring of wildlife host movement and their biotic and abiotic interactions coupled with ongoing vector, animal, and human RRV surveillance. Nevertheless, the contextualization of RRV epidemiology using macroecology may suggest potential One Health initiatives that simultaneously integrate wildlife conservation with human public health for the maximal benefit of both.

## Supporting information

## Declarations

### List of Abbreviations

RRV: Ross River virus
PSEM: Piecewise structural equation model
AIC: Akaike information criterion
PGLM: Phylogenetic generalized linear model

## Ethics approval and consent to participate

Not Applicable

## Consent for publication

Not Applicable

## Availability of data and material

All data used in this manuscript are publically available at the websites and resources described in the Methods section and in listed in the references.

## Authors’ contributions

Michael Walsh – Research conceptualization, analysis, manuscript writing, editing of drafts The author read and approved the final version of the manuscript

## Acknowledgements

Not Applicable

---

Additional file 2 Figure 1: Boxplots of species traits by taxonomic Order.

## References

1. Knope KE, Kurucz N, Doggett SL, et al (2016) Arboviral diseases and malaria in Australia, 2012–13: annual report of the national arbovirus and malaria advisory committee

2. NNDSS 2010 Annual Report Writing Group (2012) Australia’s notifiable diseases status, 2010: annual report of the national notifiable diseases surveillance system - results: zoonoses. Commun Dis Intell 36:

3. Claflin SB, Webb CE (2015) Ross River virus: many vectors and unusual hosts make for an unpredictable pathogen. PLoS Pathog 11:1–5. doi: 10.1371/journal.ppat.1005070

4. Walsh MG, Webb C (2018) Hydrological features and the ecological niches of mammalian hosts delineate elevated risk for Ross River virus epidemics in anthropogenic landscapes in Australia. Parasites Vectors 2018 111 11:192. doi: 10.1186/s13071-018-2776-x

5. Stephenson EB, Peel AJ, Reid SA, et al (2018) The non-human reservoirs of Ross River virus: a systematic review of the evidence. Parasites Vectors 2018 111 11:188. doi: 10.1186/s13071-018-2733-8

6. Russell RC (2002) Ross River virus: ecology and distribution. Annu Rev Entomol 47:1–31

7. Doherty R., Standfast H., Domrow R, et al (1971) Studies of the epidemiology of arthropod-borne virus infections at Mitchell River Mission, Cape York Peninsula, North Queensland IV. Arbovirus Infections of Mosquitoes and Mammals, 1967-1969. Trans R Soc Trop Med Hyg 65:504–513

8. Potter A, Johansen CA, Fenwick S, et al (2014) The seroprevalence and factors associated with Ross River virus infection in western grey kangaroos (Macropus fuliginosus) in Western Australia. Vector-Borne Zoonotic Dis 14:740–745. doi: 10.1089/vbz.2014.1617

9. Kay BH, Boyd AM, Ryan PA, Hall RA (2007) Mosquito feeding patterns and natural infection of vertebrates with Ross River and Barmah Forest viruses in Brisbane, Australia. Am J Trop Med Hyg 76:417–423

10. Boyd AM, Hall RA, Gemmell RT, Kay BH (2001) Experimental infection of Australian brushtail possums, Trichosurus vulpecula (Phalangeridae: Marsupialia), with Ross River and Barmah Forest viruses by use of a natural mosquito vector system. Am J Trop Med Hyg 65:777–782

11. Vale T, Spratt D, Cloonan M (1991) Serological evidence of arbovirus infection in native and domesticated mammals on the south coast of New South Wales. Aust J Zool 39:1. doi: 10.1071/ZO9910001

12. Ryan PA, Martin L, Mackenzie JS, Kay BH (1997) Investigation of gray-headed flying foxes (Pteropus poliocephalus) (Megachiroptera: Pteropodidae) and mosquitoes in the ecology of Ross River virus in Australia. Am J Trop Med Hyg 57:476–82

13. Gard G, Marshall ID, Woodroofe GM (1973) Annually recurrent epidemic polyarthritis and Ross River virus activity in a coastal area of New South Wales. II. Mosquitoes, viruses, and wildlife. Am J Trop Med Hyg 22:551–60

14. Whitehead RH (1969) Experimental infection of vertebrates with Ross River and Sindbis viruses, two group A arboviruses isolated in Australia. Aust J Exp Biol Med Sci 47:11–5

15. Lloyd-Smith JO, George D, Pepin KM, et al (2009) Epidemic Dynamics at the Human-Animal Interface. Science (80-) 326:1362–1367. doi: 10.1126/science.1177345

16. Keast A (1995) Habitat loss and species loss: the birds of Sydney 50 years ago and now. Aust Zool 30:3–25. doi: 10.7882/AZ.1995.002

17. Plourde BT, Burgess TL, Eskew EA, et al (2017) Are disease reservoirs special? Taxonomic and life history characteristics. PLoS One 12:e0180716. doi: 10.1371/journal.pone.0180716

18. Molles M (2015) Ecology[]: concepts and applications., 7th ed. Mcgraw-Hill Education, New York 19.

19. Brunner JL, LoGiudice K, Ostfeld RS (2008) Estimating reservoir competence of Borrelia burgdorferi hosts: prevalence and infectivity, sensitivity, and specificity. J Med Entomol 45:139–47

20. Ostfeld RS, Levi T, Jolles AE, et al (2014) Life History and Demographic Drivers of Reservoir Competence for Three Tick-Borne Zoonotic Pathogens. PLoS One 9:e107387. doi: 10.1371/journal.pone.0107387

21. Huang ZYX, de Boer WF, van Langevelde F, et al (2013) Species’ Life-History Traits Explain Interspecific Variation in Reservoir Competence: A Possible Mechanism Underlying the Dilution Effect. PLoS One 8:e54341. doi: 10.1371/journal.pone.0054341

22. Johnson PTJ, Rohr JR, Hoverman JT, et al (2012) Living fast and dying of infection: host life history drives interspecific variation in infection and disease risk. Ecol Lett 15:235–242. doi: 10.1111/j.1461-0248.2011.01730.x

23. Doherty RL, Gorman BM, Whitehead RH, Carley JG (1966) Studies of arthropod-borne virus infections in Queensland. V. Survey of antibodies to group A arboviruses in man and other animals. Aust J Exp Biol Med Sci 44:365–77

24. Marshall ID, Woodroofe GM, Gard GP (1980) Arboviruses of coastal south-eastern Australia. Aust J Exp Biol Med Sci 58:91–102

25. Pacioni C, Johansen CA, Mahony TJ, et al (2013) A virological investigation into declining woylie populations. Aust J Zool 61:446. doi: 10.1071/ZO13077

26. Reiss A, Jackson B, Gillespie G, et al (2015) Investigation of potential diseases associated with Northern Territory mammal declines. Charles Darwin University

27. Old JM, Deane EM (2005) Antibodies to the Ross River virus in captive marsupials in urban areas of eastern New South Wales, Australia. J Wildl Dis 41:611–614. doi: 10.7589/0090-3558-41.3.611

28. Jones KE, Bielby J, Cardillo M, et al (2009) PanTHERIA: a species-level database of life history, ecology, and geography of extant and recently extinct mammals. Ecology 90:2648–2648. doi: 10.1890/08-1494.1

29. Bielby J, Mace GM, Bininda-Emonds ORP, et al (2007) The fast-slow continuum in mammalian life history: an empirical reevaluation. Am Nat 169:748–57. doi: 10.1086/516847

30. Stekhoven DJ, Buhlmann P (2012) MissForest--non-parametric missing value imputation for mixed-type data. Bioinformatics 28:112–118. doi: 10.1093/bioinformatics/btr597

31. Liaw A, Wiener M (2002) Classification and Regression by randomForest. R News 2:18–22. doi: 10.1159/000323281

32. Shipley B (2009) Confirmatory path analysis in a generalized multilevel context. Ecology 90:363–368. doi: 10.1890/08-1034.1

33. Nunn CL, Altizer S, Jones KE, Sechrest W (2003) Comparative tests of parasite species richness in primates. Am Nat 162:597–614. doi: 10.1086/378721

34. Gómez JM, Nunn CL, Verdú M (2013) Centrality in primate-parasite networks reveals the potential for the transmission of emerging infectious diseases to humans. Proc Natl Acad Sci U S A 110:7738–41. doi: 10.1073/pnas.1220716110

35. Shipley B (2013) The AIC model selection method applied to path analytic models compared using a d-separation test. Ecology 94:560–564. doi: 10.1890/12-0976.1

36. Lefcheck JS (2016) piecewiseSEM[]: Piecewise structural equation modelling in r for ecology, evolution, and systematics. Methods Ecol Evol 7:573–579. doi: 10.1111/2041-210X.12512

37. R Core Team (2016) R: A language and environment for statistical computing. R Foundation for Statistical Computing, Vienna

38. Paradis E, Claude J, Strimmer K (2004) APE: Analyses of Phylogenetics and Evolution in R language. Bioinformatics 20:289–290. doi: 10.1093/bioinformatics/btg412

39. Popescu A-A, Huber KT, Paradis E (2012) ape 3.0: New tools for distance-based phylogenetics and evolutionary analysis in R. Bioinformatics 28:1536–1537. doi: 10.1093/bioinformatics/bts184

40. Tung Ho L si, Ané C (2014) A Linear-Time Algorithm for Gaussian and Non-Gaussian Trait Evolution Models. Syst Biol 63:397–408. doi: 10.1093/sysbio/syu005

41. Lyimo IN, Ferguson HM (2009) Ecological and evolutionary determinants of host species choice in mosquito vectors. Trends Parasitol 25:189–196. doi: 10.1016/j.pt.2009.01.005

42. Swihart RK, Gehring TM, Kolozsvary MB, Nupp TE (2003) Responses of “resistant” vertebrates to habitat loss and fragmentation: the importance of niche breadth and range boundaries. Divers Distrib 9:1–18. doi: 10.1046/j.1472-4642.2003.00158.x

43. Kosydar AJ (2014) CAN LIFE HISTORIES PREDICT THE EFFECTS O F HABITAT FRAGMENTATION? A META - ANALYSIS WITH TERRESTRIAL MAMMALS. Appl Ecol Environ Res 12:505–521. doi: 10.15666/aeer/1202_505521

44. Peñaranda DA, Simonetti JA (2015) Predicting and setting conservation priorities for Bolivian mammals based on biological correlates of the risk of decline. Conserv Biol 29:834–843. doi: 10.1111/cobi.12453

45. Dobson A (2004) Population dynamics of pathogens with multiple host species. Am Nat 164 Suppl 5:S64–78. doi: 10.1086/424681

46. Thrall PH, Antonovics J, Hall DW (1993) Host and Pathogen Coexistence in Sexually Transmitted and Vector-Borne Diseases Characterized by Frequency-Dependent Disease Transmission. Am Nat 142:543–552. doi: 10.1086/285554

47. Ferrari MJ, Perkins SE, Pomeroy LW, Bjørnstad ON (2011) Pathogens, social networks, and the paradox of transmission scaling. Interdiscip Perspect Infect Dis 2011:267049. doi: 10.1155/2011/267049

48. Fa JE, Purvis A (1997) Body Size, Diet and Population Density in Afrotropical Forest Mammals: A Comparison with Neotropical Species. J Anim Ecol 66:98. doi: 10.2307/5968

49. Walsh MG, Mor SM (2018) Interspecific network centrality, host range and early-life development are associated with wildlife hosts of Rift Valley fever virus. Transbound Emerg Dis. doi: 10.1111/tbed.12903

50. Lee KA (2006) Linking immune defenses and life history at the levels of the individual and the species. Integr Comp Biol 46:1000–15. doi: 10.1093/icb/icl049

51. Previtali MA, Ostfeld RS, Keesing F, et al (2012) Relationship between pace of life and immune responses in wild rodents. Oikos 121:1483–1492. doi: 10.1111/j.1600-0706.2012.020215.x

52. Kelly-Hope LA, Purdie DM, Kay BH (2004) Ross River virus disease in Australia, 1886-1998, with analysis of risk factors associated with outbreaks. J Med Entomol 41:133–50

53. Jacups SP, Whelan PI, Currie BJ (2008) Ross River Virus and Barmah Forest Virus Infections: A Review of History, Ecology, and Predictive Models, with Implications for Tropical Northern Australia. Vector-Borne Zoonotic Dis 8:283–298. doi: 10.1089/vbz.2007.0152

54. Flies EJ, Weinstein P, Anderson SJ, et al (2018) Ross River Virus and the Necessity of Multiscale, Eco-epidemiological Analyses. J Infect Dis 217:807–815. doi: 10.1093/infdis/jix615

55. Stratton MD, Ehrlich HY, Mor SM, Naumova EN (2017) A comparative analysis of three vector-borne diseases across Australia using seasonal and meteorological models. Sci Rep 7:40186. doi: 10.1038/srep40186

56. Shocket MS, Ryan SJ, Mordecai EA (2018) Temperature explains broad patterns of Ross River virus transmission. Elife 7:. doi: 10.7554/eLife.37762

