## Supplementary figures and images for "Ecological and life history traits are associated with Ross River virus infection among sylvatic mammals in Australia"

### Supplementary file 2

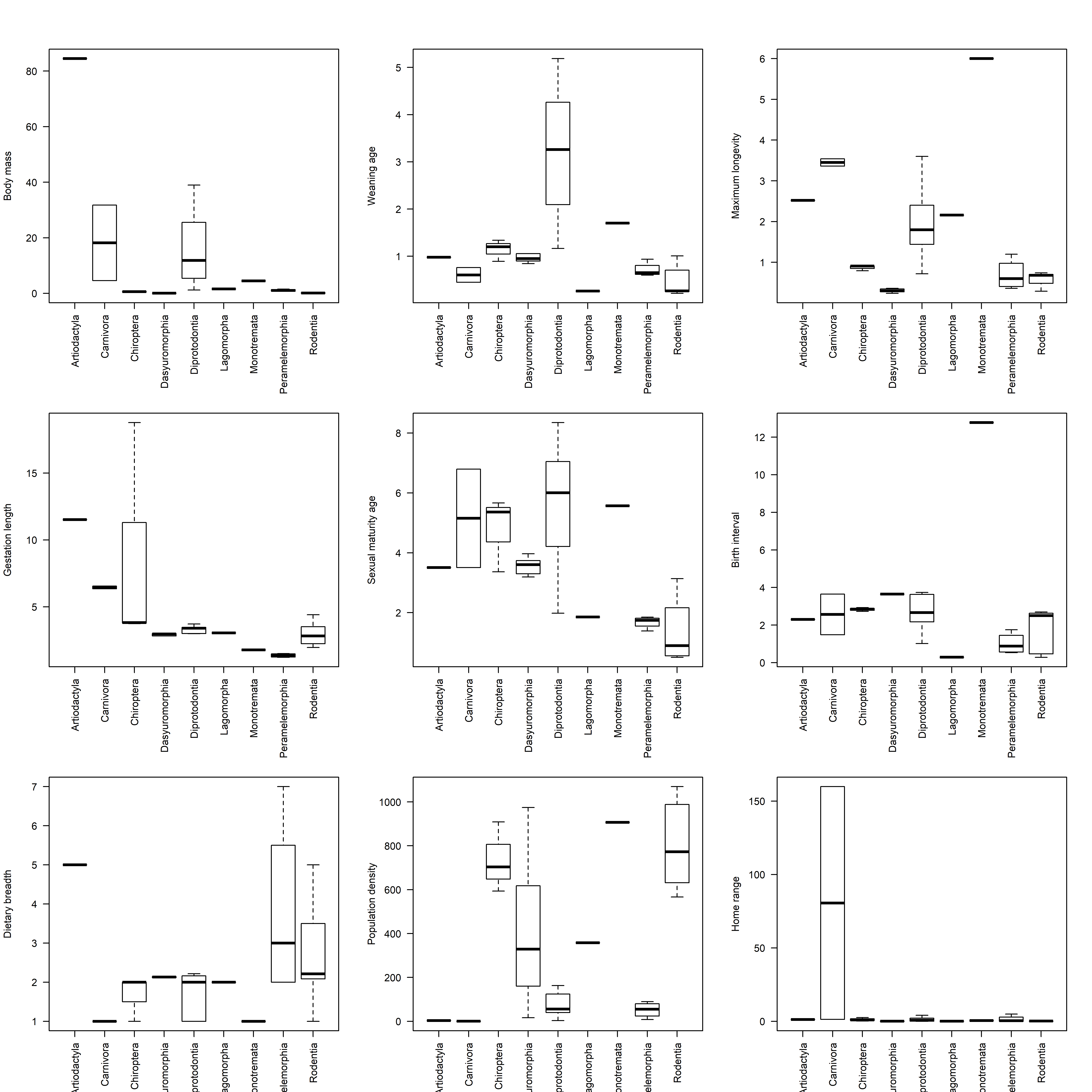
