## Supplementary material for "Ecological and life history traits are associated with Ross River virus infection among sylvatic mammals in Australia"

Additional file 2. Bivariate phylogenetic generalized linear models of the crude associations between Ross River virus infection history and biological and life history traits among sylvan Australian mammals. Each variable in the table represents a separate simple bivariate PGLM model assessing only the association between that variable alone and RRV infection history.

| Species trait | Akaike information criterion | p-value |
| --- | --- | --- |
| Diet Breadth | 94.64 | 0.0005 |
| Body mass (kg) | 107.0 | 0.39 |
| Population density (animals/km^2^) | 103.29 | 0.04 |
| Home range (km^2^) | 107.7 | 0.98 |
| Gestation length (months) | 104.17 | 0.04 |
| Litter size (animals born per litter) | 102.16 | 0.02 |
| Sexual maturity age | 105.8 | 0.18 |
| Interbirth interval | 107.0 | 0.36 |
| Weaning age | 107.7 | 0.88 |
| Maximum longevity | 107.6 | 0.70 |
